# Antimicrobial activity of polymyxin A, and characterisation of the cognate biosynthetic gene cluster within the genome of the producing *Paenibacillus polymyxa*

**DOI:** 10.64898/2026.05.29.728749

**Authors:** Amy McLeman, Alexander D. H. Kingdon, Robin Hoeven, George Taylor, Ellie Allman, Issra Bulgasim, Claudia McKeown, Richard N. Goodman, Sabrina Moyo, Adam P. Roberts

## Abstract

We report the isolation and identification of a *Paenibacillus polymyxa* strain from the citizen science project; Swab and Send. Through whole genome sequencing we are able to describe the biosynthetic gene cluster of polymyxin A produced by *P. polymyxa* 1G (NCBI accession no. JBVPZV000000000), compare the *pmxA, pmxB* and *pmxE* genes to five other polymyxin genes encoding known polymyxin variants, and provide mass spectrometry data that supports the production of polymyxin A_1_ (1157 m/z) and A_2_ (1143 m/z). Polymyxins are ranked in the highest priority critically important antimicrobials classification by the WHO and are of particular importance for treating gram-negative multidrug resistant pathogens. Due to the discovery of polymyxins occurring in the 1940’s, there is little genetic research around polymyxins, and the literature focusses primarily on clinically used polymyxin E (colistin) and polymyxin B. Previous literature suggests that polymyxin A_1_ has similar/lower toxicity to clinically used polymyxins E and B. To test if polymyxin A was able to overcome current resistance mechanisms to clinically used polymyxins, the cell free supernatant from *P. polymyxa* 1G was tested against a panel of clinical isolates with various resistance genes. We found that resistance genes *mcr-1* and *mcr-4* confer resistance to polymyxin A produced by our isolate meaning that, while polymyxin A has good antimicrobial activity, clinical resistance mechanisms already confer resistance to this variant of polymyxin.

## Introduction

Antimicrobial resistance (AMR) is one of the largest threats to global health with approximately 4.95 million associated deaths in 2019, and modelling forecasts estimating up to 8.22 million deaths associated with AMR by 2050 [2, 3]. One approach for tackling the threat of AMR is the discovery of novel antimicrobials [4-6]. Most antibiotics are derived from, or inspired by, natural products originating from bacteria or fungi, often found in soil [6, 8, 9]. Multi-drug-resistant (MDR) gram-negative bacteria are of particular concern with ever-rising numbers of infections reported; antibiotic resistant *Klebsiella pneumoniae, Acinetobacter baumannii, Pseudomonas aeruginosa* and *Escherichia coli* are some of the noted pathogens posing an important challenge in global healthcare [3, 7, 10-12]. There is potential to substantially alleviate future morbidity caused by MDR gram-negative bacteria by 11.1 million between 2025 – 2050 through the development of new antimicrobials [3].

One of the early discoveries made in the “Golden Age” of natural product discovery in 1947, by Y. Koyama, was polymyxin E, more commonly referred to as colistin [13]. *Paenibacillus polymyxa*, previously referred to as *Bacillus polymyxa*, has been extensively studied due its production of many active metabolites including multiple antibiotics such as fusaricidin and polymyxin [14, 15]. The discovery of polymyxin from *B. polymyxa* by Benedict and Langlykke (1947), and subsequent research around the time of the discovery, showed proof of antimicrobial discovery by analysing cell-free supernatant (CFS) [15, 16]. Despite how long the polymyxins have been known, there are knowledge gaps around the production of these antimicrobials, in particular genome sequence data for polymyxin biosynthesis [17]. To the best of our knowledge only five polymyxin biosynthetic gene clusters (BGCs) with the cognate polymyxin variant have been reported [18-21].

Initially thought of as a promising new antibiotic due to low levels of resistance and good activity against gram-negative bacteria, colistin (polymyxin E) was abandoned for clinical use by the 1980’s due to its toxicity [13, 18, 22, 23]. Polymyxins are associated with nephrotoxicity, neurotoxicity, allergic and topical reactions, as well as pain, fever, and even mortality [18, 22, 24]. The exact risk of incidence of nephrotoxic effects on patients receiving polymyxin treatments are highly variable across studies from 20 to 76% of patients, while meta-analysis of two large studies suggested colistin did not show significantly more cases of nephrotoxicity than other antimicrobials [22]. Older literature suggests severe events of neurotoxicity, usually associated with patients with cystic fibrosis, while newer reports suggests no or low cases of neurotoxic effects [22].

While these safety concerns surrounding polymyxins led to their discontinued clinical use, in favour of less toxic antimicrobials in the 1970’s, they have made a return to clinical use for treating MDR, particularly carbapenem-resistant, gram-negative infections [17, 25-28]. Polymyxin B and E (colistin) are the polymyxin analogues used in clinical practice and are used as a ‘last-line’ of defence for MDR gram-negative infections and they are present in the WHO’s list of medically important antimicrobials under a “sole, or one of limited available therapies” for certain infections in people [7, 17, 26, 28-30]. As such, research into polymyxin analogues with reduced toxicity is underway [7]. While polymyxin B and E have been used clinically, at least 15 naturally occurring unique polymyxins have been identified [18]. There is limited literature focussing on polymyxin A. The first record of a BGC encoding this analogue was reported in 2009 and highlighted antimicrobial activity showing polymyxin A_2_ was slightly more active than polymyxin E or B against MDR clinical isolates, and less toxic to immune cells (THP-1) [18, 31]. Polymyxin A_2_ was also less toxic to kidney cells (HEK293) than polymyxin B but showed similar toxicity to polymyxin E [31]. Most important to note here is the significant difference in effects of polymyxin A_2_ versus polymyxin A_1_, as polymyxin A_1_ showed the highest toxicity to HEK293 cells [31]. The difference between A_1_ and A_2_ is the fatty acid chain shortening from a 6-methyloctanoic acid to a 6-methylheptanoic acid [31].

Despite toxicity concerns, increased cases of MDR infections have required the use of colistin as a last line of defence [17, 23, 29, 32, 33]. Due to heavy usage in agriculture since the 1950s, antimicrobial resistance to polymyxins is widespread and of serious concern [17, 23, 29, 32, 34]. Resistance to colistin is most common through chromosomal mutations modifying the bacterial outer membrane, the lipopolysaccharide (LPS) layer, and in particular modifications of lipid A [29, 32, 35-37]. Mutations in *pmrAB, prmC, pmrE, mgrB, pmrHFIJKLM* and *PhoPQ* are reported to cause resistance to polymyxins by modifying lipid A or reducing the negative charge of the LPS which alters the way colistin interacts with the bacterial outer membrane [29, 35]. Efflux pumps are also recognised for the role they play in colistin resistance for example the *acrAB* and *kpnE* activates efflux pump systems leading to colistin resistance [29, 35].

A particular concern for colistin resistance is that of mobile resistance genes, the *mcr* genes are carried on plasmids and spread through bacterial populations via horizontal gene transfer [29, 34, 37,-39]. This means these resistance genes can be spread quickly, over wide geographical areas and across species, making them a serious concern for eliminating the effectiveness of colistin as a last resort treatment [29, 34, 37-39]. The *mcr* genes work by reducing the negative charge of the LPS through modification of lipid A [34, 37]. The first *mcr* gene (*mcr-1*) was reported in 2016 from *E. coli* isolated from a pig in China, since this, a total of 10 variants of *mcr* genes have been identified across animals, human and the environment with widespread geographical locations [29, 34-37, 40].

In this work we report on the isolation, genome sequencing and characterisation of the biosynthetic gene cluster and cognate natural product (polymyxin A) from a strain of *P. polymyxa* and determine antimicrobial activity against a range of Enterobacterales containing a diverse array of resistance genes.

## Methods

### Bacterial growth conditions

All bacterial isolates were grown on brain-heart infusion (BHI) agar at 37°C for 18 to 24 hours before growth in liquid media. Our *Paenibacillus polymyxa* 1G strain (NCBI accession number JBVPZV000000000) was grown in brain-heart infusion (BHI) or M9 glucose media at room temperature. For 500 mL M9 glucose it contained: 1x of the M9 salts in 0.5 L of water plus 500 µL of MgSO_4_ (1 M), 500 µL CaCl_2_ (0.1 M) and 8 mL of 40% (w/v) glucose. *P. polymyxa* 1G was grown at room temperature, while the resistance panel strains were incubated at 37 °C. All growth curves were performed in a CLARIOstar plate reader (BMG Labtech) at 37 °C with orbital shaking at 200 rpm between readings, for 24-hours, reading at OD_600_ using 100 flash every 10 minutes.

### Resistance panel for antimicrobial susceptibility testing

Eight clinical isolates were selected due to their differing resistance mechanisms for testing, alongside a pan-susceptible *E. coli* NCTC 86. Two of the resistance panel isolates were an *E. coli* containing *mcr-1* (RP 2) and another *E. coli* isolate containing *mcr-4* (RP 3) [41].Isolates will be referred to by their strain number as listed in table 1, with the associated resistance mechanisms described. Bioinformatic analysis was run on the whole genome sequences (WGS’s) to establish their resistance mechanisms. WGS’s were run through ResFinder with the following settings (Chromosomal point mutations: threshold for ID: 90.0%, minimum length: 60.0%, show unknown mutations: False, ignore premature stop codons: True, ignore frameshift indels: True; Acquired antimicrobial resistance genes: threshold for ID: 90.0%, minimum length: 60.0%,and specified Species and input data type as either *E. coli* or *Klebsiella*. Database versions: ResFinder-2.5., PointFinder-4.1.1) [42, 43].

**Table 1.** Strains used in the study and their resistance genes, including *E. coli* NCTC 86 strain and eight clinical isolates. Strains 1 to 7 are *E. coli* and strains 8 and 9 are *Klebsiella* spp. Strains in green grew in the presence of CFS from *P. polymyxa* 1G, while all other strains did not.

| Strain identifier | Resistance genes |  |  |  |  |  |  |
| --- | --- | --- | --- | --- | --- | --- | --- |
|  | Colistin resistance | Beta lactam | Aminoglycosides | Quinolones | Cotrimoxazole | Others | Resistance conferred by chromosomal mutations |
| ***E. coli* NCTC 86** |  | - | - | - | - | - | - |
| **RP 2** | *mcr-1.1* | *bla*CMY-2<br>*bla*TEM-1A | *ant*(3'')-Ia | - | *dfrA1*<br>*sul2* | *inu*(F) | - |
| **RP 3** | *mcr-4.6* | *bla*TEM-1A,<br>*tet*(M)<br>*tet*(A) | *ant*(3'')-Ia | - | - |  | - |
| **RP 4** |  | *bla*TEM-1B | *aph*(3'')-Ib<br>*aph*(6)-Id | - | *sul2*<br>*dfrA8* | *mph*(A) | - |
| **RP 5** |  | *bla*TEM-1B<br>*bla*CTX-M-15 | *aadA5*<br>*aph*(3'')-Ib<br>*aph*(6)-Id | - | *sul2*<br>*dfrA17*<br>*sul1* | *mph*(A) | Nalidixic Acid<br>Ciprofloxacin |
| **RP 6** |  | *bla*TEM-1B | *aph*(3'')-Ib<br>*aph*(6)-Id | - | *sul2*<br>*dfrA8* | - | - |
| **RP 7** |  | - | - | - | *sul1*<br>*dfrA7* | - | - |
| **RP 8** |  | *bla*SHV-164<br>*bla*SHV-59<br>*bla*CTX-M-15<br>*bla*TEM-1B | *aac*(3)-IId | *oqx*A<br>*oqx*B | *sul2*<br>*dfrA30* | *fosA* | Fluoroquinolone<br>Cephalosporins<br>Carbapenem |
| **RP 9** |  | *bla*SHV-110<br>*bla*SHV-81<br>*bla*CTX-M-15<br>*bla*TEM-1B | *aac*(3)-IId | *oqx*A<br>*oqx*B | *sul2*<br>*dfrA30* | *fosA6* | Fluoroquinolone<br>Cephalosporins<br>Carbapenem<br>Tigecycline |

### On-agar inhibition assay

*P. polymyxa* 1G was tested for antimicrobial activity against *Escherichia coli* NCTC 86. Colonies of *E. coli* were added to BHI broth and the OD_600_ corrected to 0.15. A sterile swab was used to spread the diluted *E. coli* culture across a BHI agar plate (4% (w/v) agar) to make a background lawn of microbial growth. A spot of *P. polymyxa* 1G was added onto the lawn and the agar plate was incubated (2 days, room temperature) and zone of inhibition assessed.

### Cell-free supernatant inhibition assay

After *P. polymyxa* 1G showed a zone of inhibition on agar it was screened in a CFS assay against either *E. coli* or the resistance panel strains. *P. polymyxa* 1G was grown for 18 hours (10ml BHI, 37 °C) and for 2 weeks (10 mL, M9 Glucose, room temperature). One mL aliquots of the cultures were centrifuged at 13,000 x g for 1 minute and the supernatant was filtered (0.22 µm) to produce the CFS. Each well of a 96-well plate (Costar) had either CFS added (100 µL), or control wells containing CFS from the *E. coli* or the resistance panel strains (100 µL). This was followed by addition of diluted cultures of the test strains (1:5,000 dilution of an OD_600_ = 0.1 culture of an *E. coli*, or one of the resistance panel strains, overnight cultures, all into fresh BHI). Positive control wells had 100 µL BHI and the 100 µL diluted *E. coli*, or resistance panel strain, and negative control wells had 200 µL BHI. Growth was measured for 24-hours using a CLARIOstar plate reader as described above.

The resulting data was analysed using GraphPad Prism (version 10.4.0) software to calculate the area under the curve. The reduction in area under the curve is calculated by comparing the CFS from *E. coli* or resistance panel strain respectively, with *E. coli* growth in each environmental isolates’ CFS. Growth in the presence of colistin sulphate (Apollo Scientific) was assayed using the same protocol for the resistance panel (strains 1-4), except colistin diluted in molecular water replaced CFS in each well of the 96-well plate.

### Isolate identification and sequencing

Strains were grown for two days (room temperature on BHI agar). Colony PCR used an individual colony of each isolate, MyTaq 2x Master Mix, and the following primers (27F - AGAGTTTGATCCTGGCGCAG and 1492R - GGTTACCTTGTTACGACTT) with PCR conditions: 95 °C, 10 minutes; 35 cycles (95 °C, 1 min; 50 °C, 30 sec; 72 °C, 30 sec) and a final 72 °C, 5 min. PCR cleanup was performed using the Monarch^®^ Spin PCR & DNA Cleanup Kit (New England Biolabs). The purified DNA was then sent to GeneWiz for Sanger sequencing. The resulting sequence was matched to other 16S rRNA sequences using BLASTn against the core nucleotide database, to identify the closest species matches.

*P. polymyxa* 1G was also sent for whole genome sequencing. MicrobesNG (MicrobesNG, Birmingham, UK) performed short read sequencing using HiSeq X10 (Illumina, San Diego, CA, USA). Genomic DNA was extracted using a Fire Monkey High Molecular Weight DNA Extraction Kit (RevoluGen, Berkshire, UK), and long read sequencing was carried out using Oxford Nanopore Technologies (Oxford, UK) R9.4.1 flow cell on a MinION sequencer. The reads were basecalled with MinKNOW software using Guppy algorithm (v4.0.9). Long reads were trimmed using Porechop (https://github.com/rrwick/Porechop) and filtered using Filtlong (https://github.com/rrwick/Filtlong), and the genome assembly was performed using Unicycler (v0.4.8.0) with default parameters. The assembly yielded a genome sequence of 6,088,001 bp with a G+C content of 45.7%. The genome was annotated using RAST (v2.0).

### Bioinformatics analysis of *P. polymyxa* 1G whole genome sequence

The assembled *P. polymyxa* 1G genome was run through antiSMASH (version 8.0) to look for known BGCs. The polymyxin BGC was focussed upon and the *pmxA, pmxB* and *pmxE* genes encoding the biosynthetic proteins were analysed in more detail. Available *pmx* genes associated with characterised polymyxin variants were aligned with our isolate’s genes, using Clustal Omega multiple sequence alignment [44] and the tree was created using iTOL (version 7.2). Alignment sequence Genbank accession numbers were; polymyxin A from *P. polymyxa* (EU371992.1) [18], a polymyxin B variant with D-dab at region 3 from *P. polymyxa* (JN660148.1) [20], polymyxin E from *P. alvei* (**LS**992241.1) [21], polymyxin E from *P. alvei* (**KP**262070.1) [21] and polymyxin P from *P. polymyxa* (HE577054.1) [19]. The two polymyxin Es will be differentiated using the first two letters of their accession number in the following text.

Supplementary table 2 showing the active site residues for the polymyxin variants was created by extracting active site residues from antiSMASH (v8.0) using the Stachelhaus sequence. While the stereochemical configuration of each amino acid was predicted from the antiSMASH NPRS/PKS module view, with the presence of an epimerisation domain indicating the D-stereoisomer of an amino acid.

### Mass Spectrometry

The *P. polymyxa* 1G isolate was cultured in M9 Glucose for two weeks (room temperature). The samples were mixed with acetonitrile (100 µL + 100 µL) to remove protein and centrifuged at 20,000 x *g* for 3 min at 4 °C. The supernatant was then dried under nitrogen gas flow at 40 °C and resuspended in 100 µL 5% acetonitrile in water.

Liquid chromatography-mass spectrometry (LC-MS) analysis was performed using a SCIEX Exion LC system consisting of two AD high pressure gradient pumps, vacuum degasser, solvent valve, AC column oven and AC Autosampler, coupled to a SCIEX 7600 ZenoTOF Q-TOF mass spectrometer with TurboV Optiflow ion source running a 50 µm ESI probe. The system was controlled by SCIEX OS v3.0.

A sample volume of 10 μL was injected onto a 50 μL sample loop. Injection cycle time was 1 min per sample. Separations were performed using a Thermo Accucore C18 column with dimensions of 150 mm length, 2.1 mm diameter and 2.6 μm particle size equipped with a guard column of the same phase. Mobile phase A was water with 0.1% formic acid; mobile phase B was 98% acetonitrile and 2% water with 0.1% formic acid. Separation was performed by gradient chromatography at a flow rate of 0.3 mL/min, starting at 5% B for 1 minute, ramping to 100% B over 7 min, hold at 100% B for 2 min, then back to 5% B. Re-equilibration time was 4 min. Total run time including injection cycle was 15 min.

The mass spectrometer was run in positive mode under the following source conditions: curtain gas pressure, 50 psi; ion spray voltage, 5500 V; temperature, 400 °C; ESI nebulizer gas pressure, 50 psi; heater gas pressure, 70 psi; declustering potential, 30 V. Mass spectrum data was acquired in the mass range of 120 – 1200 *m/z* with an accumulation time of 0.25 s. Declustering potential was 50 V and collision energy was 10 V. Colistin was analysed by measuring the peak areas of the [M+2H]^2+^ and [M+3H]^3+^ ions at *m/z* 578.383 and 385.924 ± 0.05 respectively, with the [M+3H]^3+^ ion acting as quantifier and the [M+2H]^2+^ ion acting as qualifier. The retention time of colistin was determined to be 3.48 min. Polymyxin A_1_ and A_2_ were identified as the [M+H]^+^ ions at m/z 1157.74 and 1143.72, eluting at 3.483 and 3.324 minutes from the *P. polymyxa* 1G CFS.

## Results

The *P. polymyxa* 1G strain was isolated from a swab of a coal scuttle in 2019 from the Swab and Send project [45]. The isolate showed a zone of inhibition against *E. coli* on agar screening and the follow up CFS assay showed a 99% reduction in area under the curve when compared to the growth of *E. coli* in spent media as shown in figure 1. The almost total inhibition of growth of *E. coli* led to species identification of the *P. polymyxa* 1G isolate, with the closest match of the 16S rRNA sequence being to *Paenibacillus polymyxa*. A simpler growth media background was required for downstream data processing using mass spectrometry, so the isolate was grown in M9 using glucose as the carbon source. The *P. polymyxa* 1G isolate was grown in M9 media for 2 weeks and retention of activity was confirmed by comparing the activity of BHI broth CFS to M9 broth CFS causing a 99% and 98% reduction in area under the curve, respectively (figure 1). The compounds produced by the *P. polymyxa* 1G strain showed a similar pattern of mass ion peaks to colistin. Colistin sulphate bought from Apollo Scientific was used as a positive polymyxin control for mass spectrometry. The comparison between mass spectrometry spectra for the two peaks within our sample and the commercial colistin sulphate sample can be seen in figure 2.

**Figure 1.**
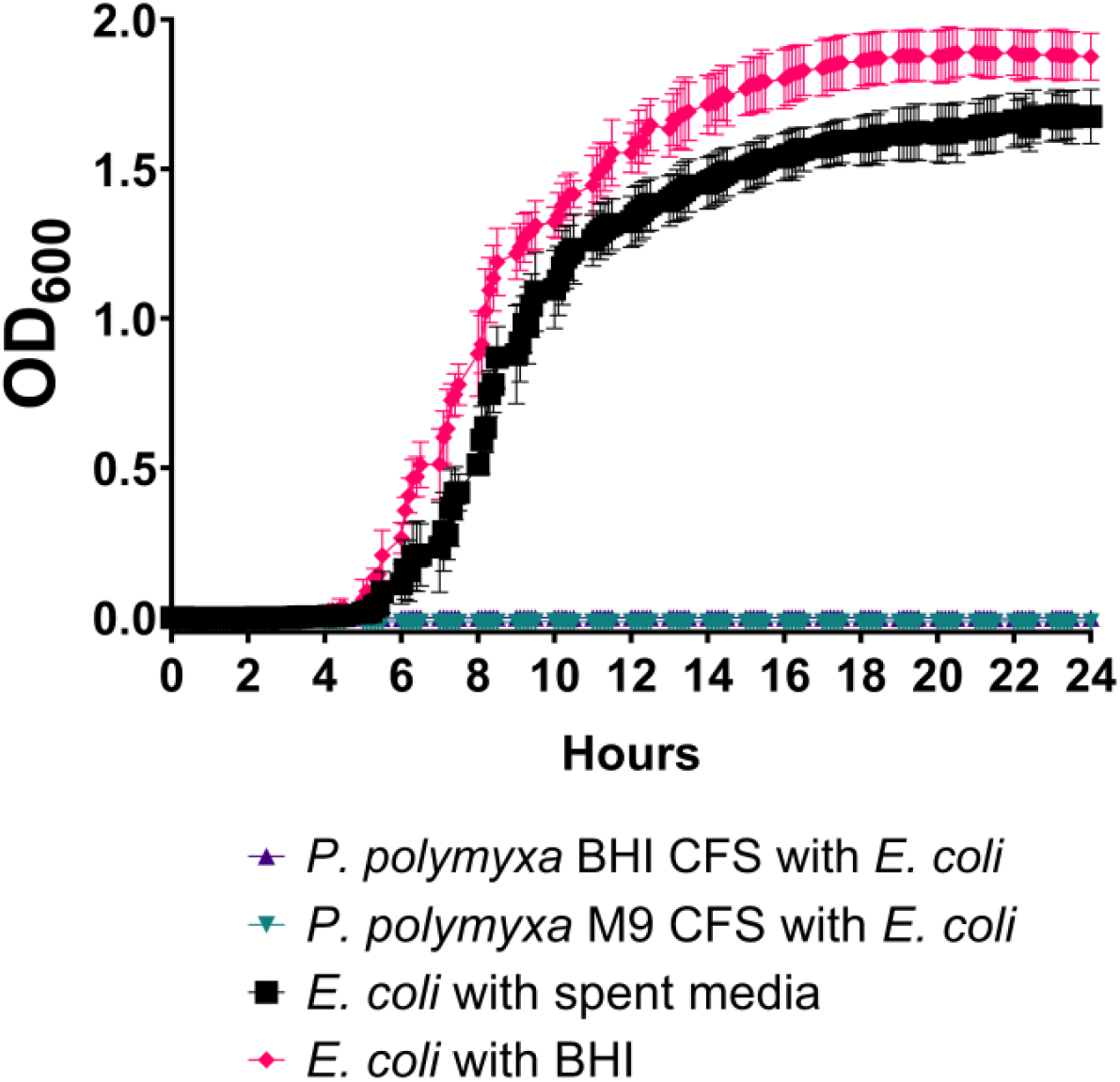
*E. coli* growth curves showing the inhibitory effects of *P. polymyxa* CFS. The growth of *E. coli* when grown in the presence of the CFS from our isolate 1G versus with BHI broth only (control 1) or with CFS from the *E. coli* overnight, spent media (control 2). Control 1 shows the normal growth of *E. coli* in nutrient rich broth, control 2 accounts for the loss of nutrients when an isolate is grown in media overnight. Our isolate 1G was grown in BHI and M9 media. OD_600_ = absorbance at 600 nm.

**Figure 2.**
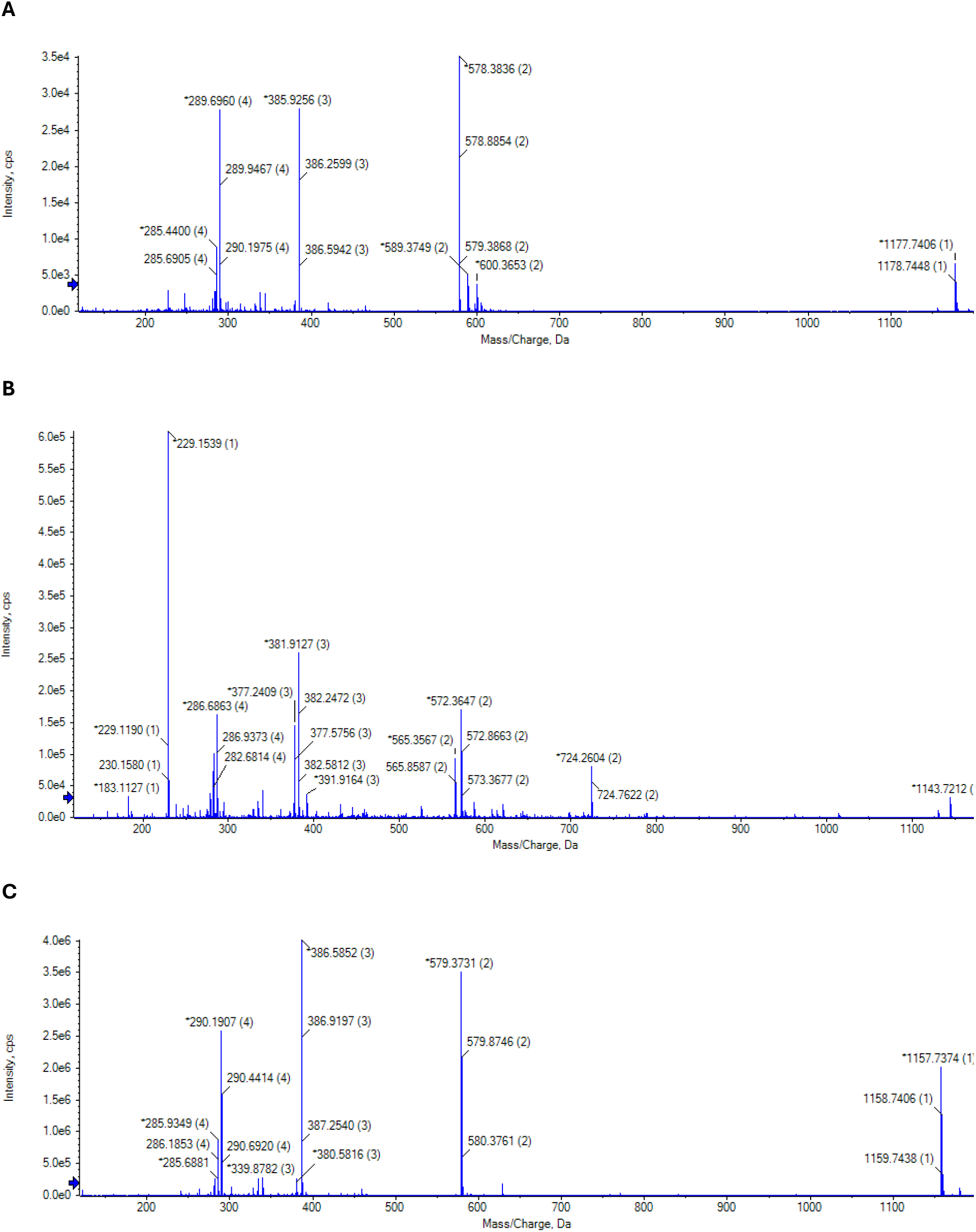
Mass spectrometry spectra summarising masses detected at specific peaks of LC-MS elution for **A** colistin sulphate eluting at 3.576 minutes, **B** *P. polymyxa* 1G peak eluting at 3.324 minutes and **C** *P. polymyxa* 1G peak eluting at 3.483 minutes.

Colistin sulphate eluted at 3.576 minutes, while our isolate had a large peak eluting at 3.483 minutes and a second peak of interest eluting at 3.324 minutes (supplementary figure 1), both of which appear to have similar ion charge M^1+^, M^2+^, M^3+^, and M^4+^ patterns to colistin in the mass spectrum (figure 2). The molecular ions from the peaks eluting at 3.483 and 3.324 minutes were 1157.74 [M+H]^+^ and 1143.72 [M+H]^+^, respectively. In our experiment, the polymyxin E (colistin) molecular ion showed an m/z value of 1177.74 [M+Na]^+^, but has also been reported as the molecular ion of 1155.74 [M+H]^+^ [46, 47]. While the ion charge patterns were similar to known polymyxin E, the difference in mass suggests our isolate is not producing polymyxin E. Literature shows a close match to our two peaks with two compounds m/z values of 1143.69 [M+H]^+^ and 1157.69 [M+H]^+^ [31]. These two compounds were identified as polymyxin A variants A_1_ and A_2_, with a difference of 14 Daltons corresponding to a methylene (CH_2_) group. Our analogues have the same difference between our two peaks m/z values, equating to a longer N-terminal fatty acid chain [31]. We therefore predict that our isolate is also producing these two variants of polymyxin A.

To qualify the prediction of the polymyxin produced by our isolate being polymyxin A, the WGS was obtained and run through antiSMASH to investigate the predicted BGCs encoded by our isolate. This resulted in a high confidence match to polymyxin, among several other high confidence matches (supplementary table 1). One output from antiSMASH is a prediction of the amino acids which are being incorporated into a final compound being produced by non-ribosomal peptide synthetases (NRPS). Each NRPS domain is responsible for addition of a single amino acid into the final compound. The predicted amino acids forming the encoded polymyxin were extracted from the antiSMASH output and compared to the expected amino acids for each of the other previously characterised polymyxin analogues (figure 3). This figure shows that the predicted polymyxin produced by our isolate matches polymyxin A. Both our encoded polymyxin and polymyxin A had the same two amino acid differences, at region 3 (R3), 6 (R6) and 7 (R7), from the other polymyxin variants (supplementary table 2).

**Figure 3.**
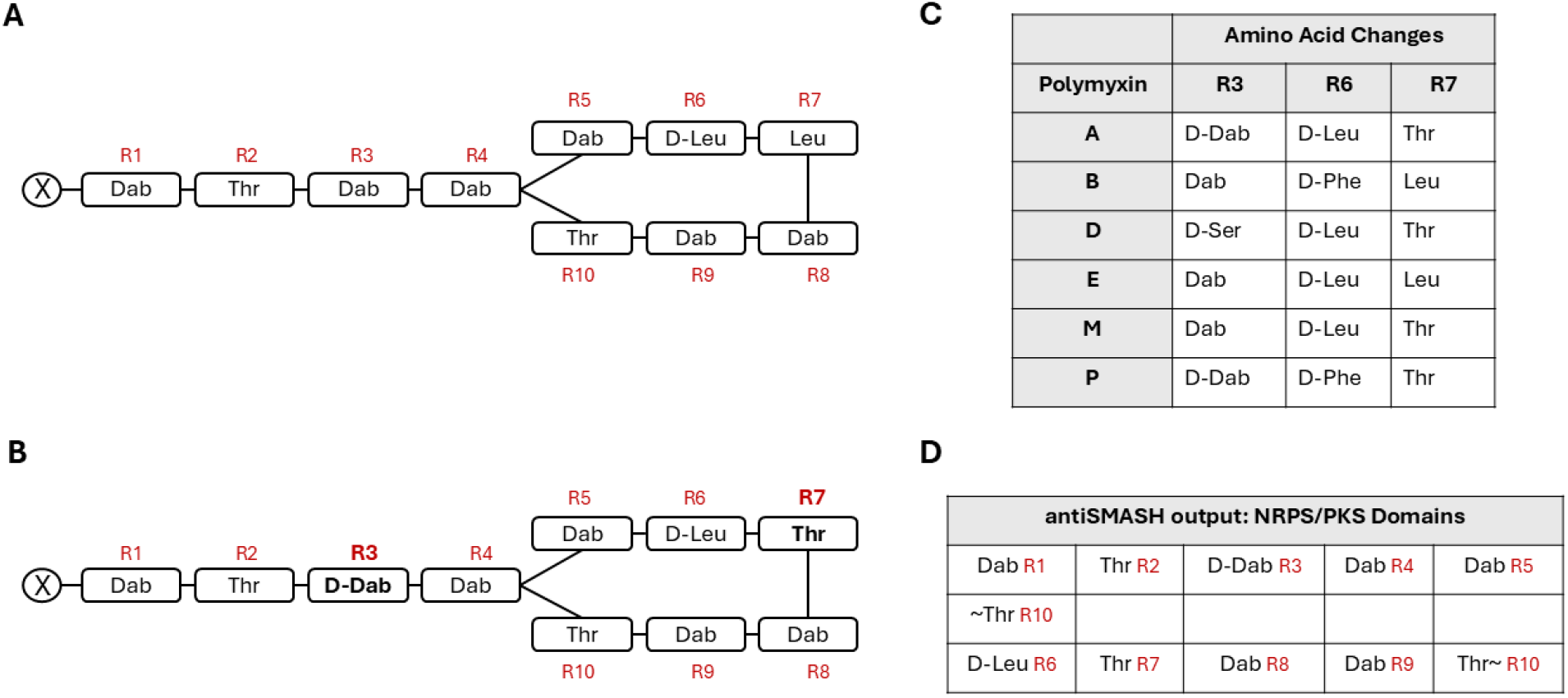
showing the amino acid structures and regions (R) of **A** colistin/polymyxin E [1], **B** the polymyxin amino acid structure of our isolate, **C** a table of the amino acid changes at the respective regions for the different polymyxin analogues (adapted from [7]) and **D** the antiSMASH output for the NPRS/PKS domains used to show the amino acid structure of our predicted compound.

Polymyxin biosynthesis is encoded for by three NRPSs (*pmxA, pmxB* and *pmxE*) and two ABC-type transporters. Each *pmx* NRPS gene from our isolate was aligned with the other polymyxin analogue *pmx* NRPS genes, using ClustalOMEGA to determine the relatedness. The phylogenetic trees of the *pmxA* genes for all polymyxin analogues are shown in figure 4, alongside the same procedure for *pmxB* and *pmxE*.

**Figure 4.**
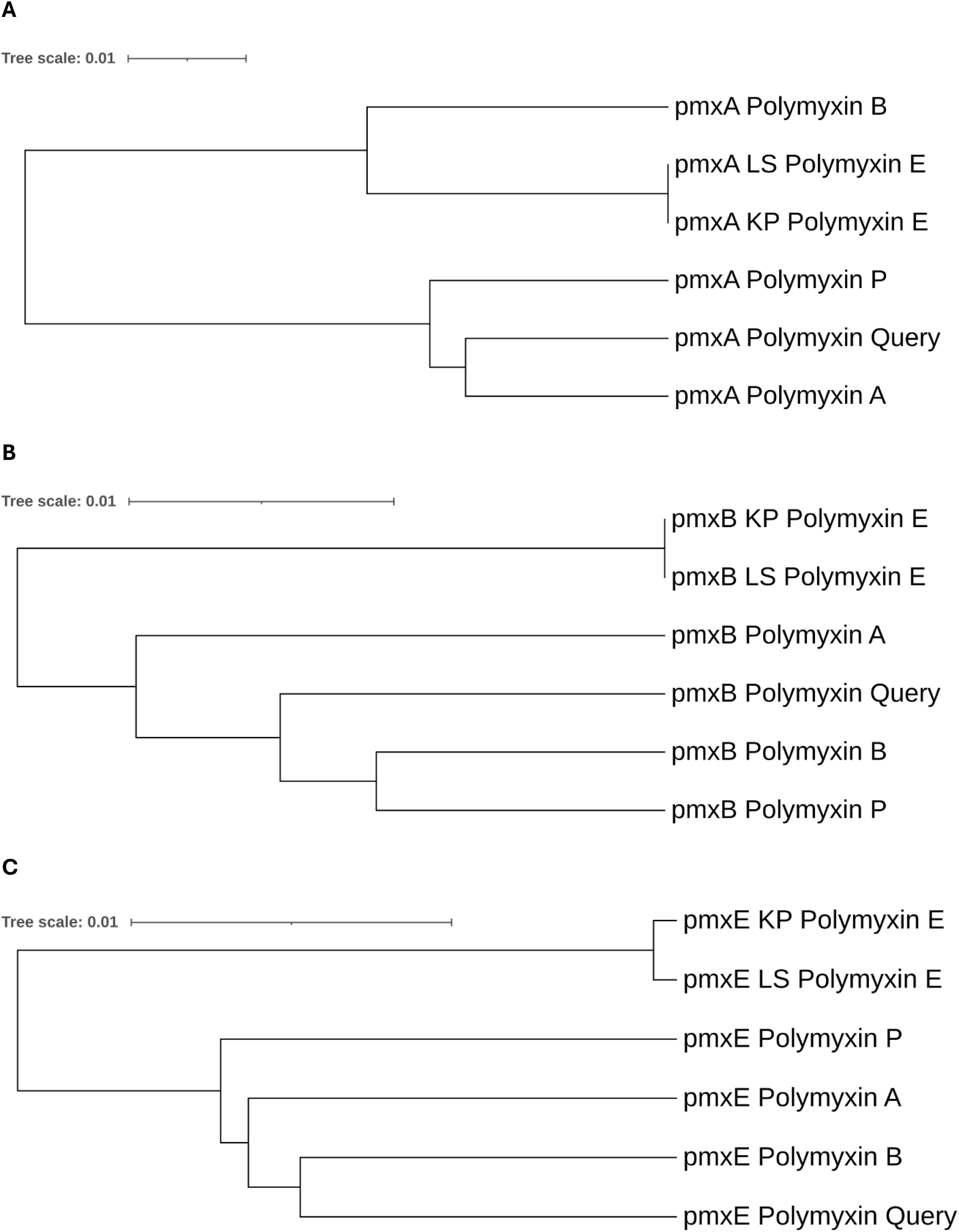
showing the guide tree alignment of the three *pmx* genes **A** *pmxA*, **B** *pmxB* and **C** *pmxE* for all polymyxin analogue protein sequences when the polymyxin analogue is also known. Important note: the polymyxin B aligned here is for a variant of polymyxin B that has a D-Dab at region 3 instead of the expected L-Dab.

The three *pmx* genes encode different regions of the polymyxin; *pmxE* encodes regions 1 to 5, *pmxA* encodes regions 6 to 9 and *pmxB* encodes region 10 (figure 3). As *pmxB* only encodes for a threonine at region 10, for all known polymyxins with associated gene sequences, significant differences in the sequence would not be expected. Although the grouping of both polymyxin E analogues would be expected as both sequences are from *P. alvei* instead of *P. polymyxa*. While *pmxE* encodes the first five regions, for the four characterised analogues, only polymyxin E differs in containing an L-Dab (2,4 -diaminobutyric acid) instead of a D-Dab in region 3, thus, we would expect clustering of the polymyxin E encoding *pmxE* genes separate from the other polymyxin analogue *pmxE* genes. An important note that while normally we would also expect this difference for polymyxin B, the characterised variant encodes a D-Dab at region 3, removing this expected difference between polymyxin B and polymyxins A and P.

The most important alignment for distinguishing the polymyxin analogues is the *pmxA* gene due to it encoding both region 6 and 7, which are the variable regions for each polymyxin analogue (see figure 3 and 4). Both *pmxA* genes encoding polymyxin E cluster together, potentially due to them being from a different *Paenibacillus* species, but also due them encoding D-leucine and L-leucine at regions 6 and 7 respectively. Closely clustering to the polymyxin E *pmxA* is polymyxin B’s *pmxA*, encoding the same amino acid at region 7 but D-phenylalanine at region 6 instead. While the polymyxin P *pmxA* gene clusters more closely with polymyxin A *pmxA* gene, this is likely due to region 7 encoding a leucine in polymyxin B and E, versus a threonine in polymyxin P and A. The higher weighting in relatedness for changes at region 7 versus region 6 is further explained by the increased variation in the active site residues of region 7 (supplementary table 2). The clustering of our query polymyxin sequence with polymyxin A and the conserved active site residues at every region between these two sequences supports the identification of *P. polymyxa* 1G’s polymyxin as polymyxin A.

We tested eight resistant isolates against the CFS of *P. polymyxa* 1G strain and found that the isolates with MCR resistance genes (RP 2 and RP 3) were resistant to polymyxin A, as expected, and isolate RP 4 was also resistant (figure 5). Resistance from isolate RP 4 was unexpected as there are no polymyxin resistance genes or chromosomal mutations, a follow up experiment was run to test resistance to colistin sulphate salt using *E. coli* NCTC 86 as a negative control and RP 2 and RP 3 as positive controls (supplementary figure 2). Both *E. coli* isolates containing MCR resistance genes are resistant to the highest tested concentration of colistin 4 mg/L surpassing the clinical breakpoint for colistin resistance at 2 mg/L. *E. coli* NCTC 86 was able to grow normally in the presence of colistin up to 0.5 mg/L while isolate RP 4 was able to grow relatively well in the presence of colistin at 1 mg/L. Susceptibility of *E. coli* strains is shown to be the same across polymyxin E and A [31]and so this data suggests that growth of RP 4 in the presence of *P. polymyxa* 1G CFS versus the lack of growth of *E. coli* NCTC 86 may be because this isolate is producing approximately 1 mg/L of polymyxin A.

**Figure 5.**
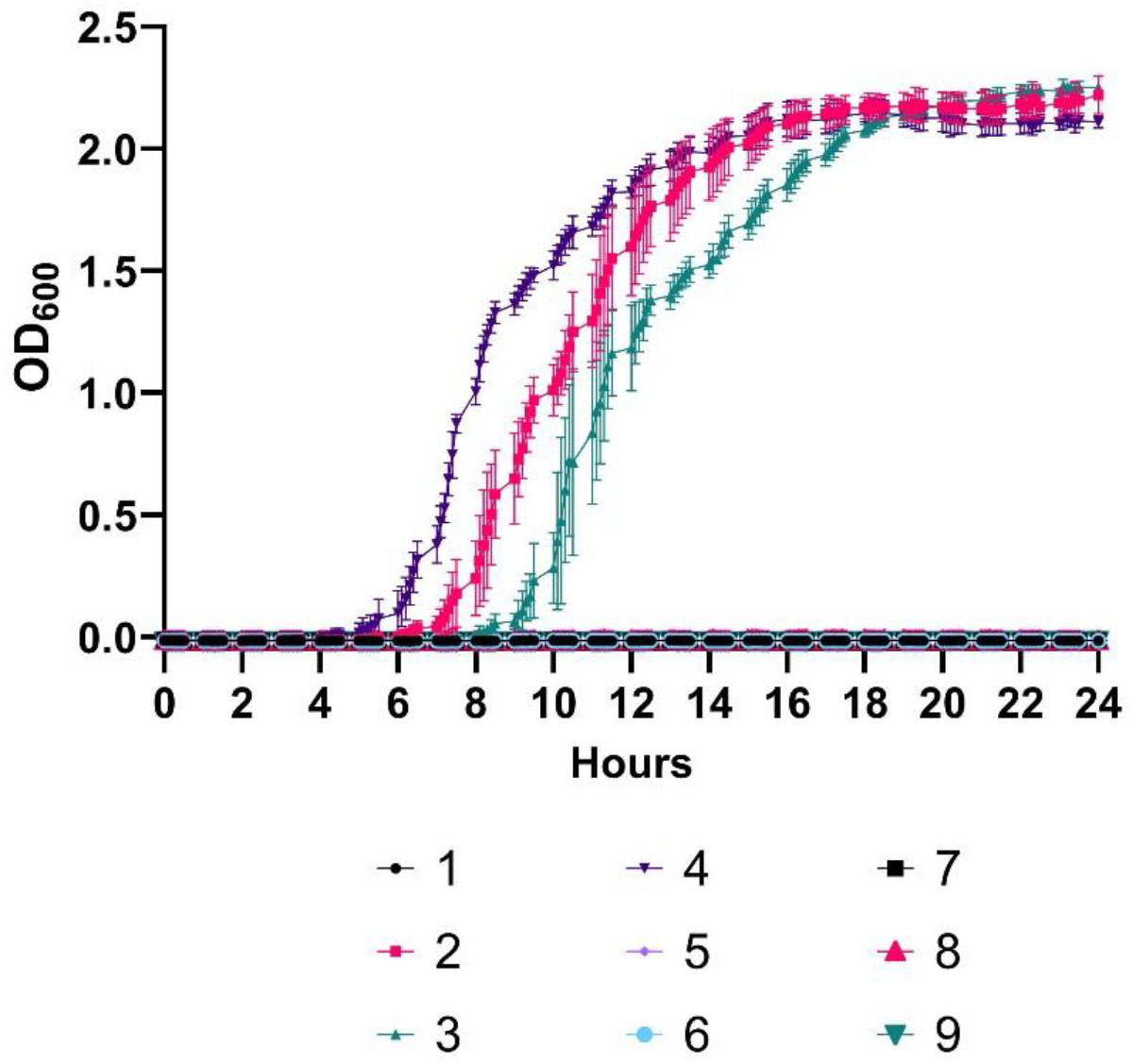
showing growth of all 9 RP isolates in the presence of CFS from *P. polymyxa* 1G grown in M9 which has mass spectrum data to support the presence of polymyxin. Here we see resistance to our CFS containing polymyxin A from isolates 2 and 3 containing MCR 1 and MCR 4 respectively. And resistance from RP isolate 4 which has supporting biological data to show low level resistance to colistin (supplementary figure 2).

## Discussion

In this study, we have shown the mass spectrum data for our isolate’s antimicrobial CFS containing two compounds with m/z values of 1143.7212 and 1157.7374, which show very similar m/z values to two polymyxin A variants characterised by Jangra *et*.*al*, 2018 of 1143.69 and 1157.69 [31]. Jangra *et al*., 2018 showed the 14 Da difference between the compounds was a difference in the N-terminal fatty acid chain length, from 6-methyloctanoic acid in polymyxin A_1_ (1157 m/z) to 6-methylheptanoic acid in polymyxin A_2_ (1143 m/z) [31]. So, rather than two distinct antimicrobial compounds being produced by *P. polymyxa* 1G, we are likely seeing the two variants of polymyxin A as Jangra *et*.*al*, 2018 described [31]. This is supported by sequence data from our isolate *P. polymyxa* 1G showing the predicted amino acid structure (figure 3) and active site residues match (supplementary table 2) between our sequence and the sequence encoding polymyxin A. We also show the alignment of our query sequence against all five polymyxin BGCs. As the differences for polymyxin A compared to polymyxin B, E and P are most significant at regions 6 and 7, encoded for by *pmxA*, the clustering of our sequence with polymyxin A encoding *pmxA* for this NRPS also validates that our isolate is producing polymyxin A.

Clinically used polymyxin variants are restricted to polymyxin E and polymyxin B due to them having the lowest nephrotoxicity amongst tested analogues in *in vivo* models in the 1960’s [23]. However, more recent research suggests that polymyxin A_2_ has lower toxicity to an immune cell line, similar toxicity to a kidney cell line and better efficacy than polymyxin B and E against MDR clinical bacterial isolates [31]. Jangra *et al*., 2018, suggested further research was need into polymyxin A’s potential clinical use, prompting us to test the polymyxin A from this *P. polymyxa* 1G strain against eight clinical isolates with various resistance gene profiles, and a pan-susceptible *E. coli* NCTC 86 strain. As polymyxin is a last line of defence antibiotic important for treating Gram-negative infections, and colistin resistance is particularly widespread in *E. coli* and *Klebsiella* spp. infections, a selection of clinical isolates were chosen to test against [38, 48, 49]. Two of the resistance panel isolates, RP 2 and RP 3, contained *mcr* genes which confer resistance to polymyxins by modifying lipid A in the outer membrane lipopolysaccharides [34]. *mcr-1*, which is present in RP 2, is globally widespread and we have shown it confers resistance to polymyxin A [34]. *mcr-4*, present in the RP 3, is a less common *mcr* gene, which we have shown can also confer resistance to polymyxin A [34, 50, 51].

Lastly, RP 4 showed resistance to the CFS of *P. polymyxa* 1G but had no known resistance genotype polymyxins. Biological data collected in this study showed over 50% of expected growth in 1 mg/L colistin compared to only 5% growth of the *E. coli* NCTC 86 indicator strain (Supplementary figure 2). Jangra *et al*., 2018, show that there is negligible, if any difference, between the MICs of different polymyxin variants A, B and E against *E. coli* and *Klebsiella* spp. strains. Combined with our data, this suggests our strain could be producing polymyxin at a concentration of approximately 1 mg/L, allowing us to see growth at just below the colistin EUCAST breakpoint of 2 mg/L in RP 4, but not in the pan-susceptible *E. coli* NCTC 86 strain. An alternative explanation is that there is a novel polymyxin resistance mechanism and further work is ongoing to investigate this possibility.

## Conclusions

*P. polymyxa* 1G identified from the Swab and Send project [45] has been shown here to produce polymyxin A with mass spectrum data, and genetic data matching the predicted amino acid structure, of polymyxin A. Sequence alignment of the *pmx* genes encoding the amino acids differences between polymyxin variants also align with polymyxin A. While previous toxicity data suggested polymyxin A may be a good alternative to polymyxin E and B, we found that clinical resistance mechanisms also confer resistance to polymyxin A. Though it may have similar or less toxicity to clinically used polymyxins, it is unlikely to be an alternative for use against pathogens resistant to the last line of defence clinical polymyxins.

## Ethics Statement

The Authors have not performed any study on human subjects or animal models.

## Author Contributions

AM, EA, IB, CM, RH and GT performed experiments. Data analysis carried out by AM, AK, GT. RG and SM provided resources. AM and AK wrote and edited the manuscript. APR supervisor, funding acquisition and conceptualisation. All authors contributed to the manuscript and approved the final version.

