## Supplemental data for "Antimicrobial activity of polymyxin A, and characterisation of the cognate biosynthetic gene cluster within the genome of the producing *Paenibacillus polymyxa*"

**Supplementary Material for Swab and Send, citizen science project, identifies isolate with antimicrobial activity against *Escherichia coli* as *Paenibacillus polymyxa* producing polymyxin A**

Amy McLeman[1], Alex D. H. Kingdon[1], Robin Hoeven[2], George Taylor[2], Ellie Allman[1], Issra Bulgasim[1], Claudia McKeown[1], Richard N. Goodman[1], Sabrina Moyo[1], and Adam P. Roberts[1,#].

**Supplementary Table 1** showing the output of antiSMASH 8.0 from running the whole genome sequence of *Paenibacilus polymyxa* 1G using relaxed detection strictness, and the extra features: Known Cluster Blast, TFBS analysis, Active Site Finder, Sub Cluster Blast and RREFinder.

| Region | Type | From | To | Similarity confidence | Most similar known cluster |
|---|---|---|---|---|---|
| **Region 1** | NRPS | 67,893 | 131,619 | **High** | fusaricidin B |
| **Region 2** | RRE-containing | 379,655 | 401,109 | | |
| **Region 3** | NI-siderophore | 1,031,093 | 1,066,494 | | |
| **Region 4** | Proteusin | 1,216,382 | 1,236,618 | | |
| **Region 5** | NRPS,transAT-PKS | 1,272,973 | 1,372,957 | | |
| **Region 6** | lassopeptide | 1,408,291 | 1,432,389 | **Low** | paeninodin |
| **Region 7** | lanthipeptide-class-i | 1,776,601 | 1,803,608 | **High** | paenilan |
| **Region 8** | NRPS-like | 2,128,959 | 2,172,654 | | |
| **Region 9** | cyclic-lactone-autoinducer | 2,183,714 | 2,204,357 | | |
| **Region 10** | NRPS | 2,549,080 | 2,642,697 | **High** | tridecaptin |
| **Region 11** | transAT-PKS,NRPS | 2,731,709 | 2,801,719 | **Low** | Pelgipeptin A/ B/ C/ D |
| **Region 12** | transAT-PKS,NRPS | 2,816,149 | 2,898,822 | **Medium** | paenilipoheptin |
| **Region 13** | cyclic-lactone-autoinducer | 3,248,592 | 3,269,140 | | |
| **Region 14** | terpene-precursor | 3,601,295 | 3,622,218 | | |
| **Region 15** | transAT-PKS,NRPS,T3PKS,PKS-like | 3,788,534 | 3,890,613 | **Low** | aurantinin B/ C/ D |
| **Region 16** | terpene-precursor | 4,243,400 | 4,265,778 | | |
| **Region 17** | NRPS | 5,216,214 | 5,297,097 | **High** | polymyxin |
| **Region 18** | phosphonate | 5,895,571 | 5,946,724 | | |

**Supplementary Table 2** showing the active site residues for the five polymyxin analogues and the polymyxin query sequence from *P. polymyxa* 1G. Highlights show the differences in active site residues relative to our isolate's Pmx proteins predicted active site residues.

| Region | Accession number | Genbank accession | Polymyxin Analogue | Important active site residues in adenylation-domain | | | | | | | | | | Specified amino acid |
|---|---|---|---|---|---|---|---|---|---|---|---|---|---|---|
| | | *Region 1* | | | | | | | | | | | | |
| **pmxE1** | Polymyxin Query | | | D | V | G | E | I | S | S | I | D | K | L-Dab |
| | E681 | EU371992.1 | Polymyxin A | D | V | G | E | I | S | S | I | D | K | L-Dab |
| | M-1 | HE577054.1 | Polymyxin P | D | V | G | E | I | S | S | I | V | K | L-Dab |
| | | KP262070.1 | Polymyxin E | D | V | G | E | I | S | S | I | D | K | L-Dab |
| | B-LR | LS992241.1 | Polymyxin E | D | V | G | E | I | S | S | I | D | K | L-Dab |
| | PKB1 | JN660148.1 | D-Dab3-Polymyxin B | D | V | W | E | I | S | S | I | D | K | L-Dab |
| | | *Region 2* | | | | | | | | | | | | |
| **pmxE2** | Polymyxin Query | | | D | F | W | N | I | G | M | V | H | K | L-Thr |
| | E681 | EU371992.1 | Polymyxin A | D | F | W | N | I | G | M | V | H | K | L-Thr |
| | M-1 | HE577054.1 | Polymyxin P | D | F | W | N | I | G | M | V | H | K | L-Thr |
| | | KP262070.1 | Polymyxin E | D | F | W | N | I | G | M | V | H | K | L-Thr |
| | B-LR | LS992241.1 | Polymyxin E | D | F | W | N | I | G | M | V | H | K | L-Thr |
| | PKB1 | JN660148.1 | D-Dab3-Polymyxin B | D | F | W | N | I | G | M | V | H | K | L-Thr |
| | | *Region 3* | | | | | | | | | | | | |
| **pmxE3** | Polymyxin Query | | | D | V | G | E | I | S | S | I | D | K | D-Dab |
| | E681 | EU371992.1 | Polymyxin A | D | V | G | E | I | S | S | I | D | K | D-Dab |
| | M-1 | HE577054.1 | Polymyxin P | D | V | G | E | I | S | S | I | D | K | D-Dab |
| | | KP262070.1 | Polymyxin E | D | V | G | E | L | S | S | I | D | K | L-Dab |
| | B-LR | LS992241.1 | Polymyxin E | D | V | G | E | L | S | S | I | D | K | L-Dab |
| | PKB1 | JN660148.1 | D-Dab3-Polymyxin B | D | V | G | E | I | S | S | I | D | K | D-Dab |
| | | *Region 4* | | | | | | | | | | | | |
| **pmxE4** | Polymyxin Query | | | D | V | G | E | I | S | A | I | D | K | L-Dab |
| | E681 | EU371992.1 | Polymyxin A | D | V | G | E | I | S | A | I | D | K | L-Dab |
| | M-1 | HE577054.1 | Polymyxin P | D | V | G | E | I | S | A | I | D | K | L-Dab |
| | | KP262070.1 | Polymyxin E | D | V | G | E | I | S | A | I | D | K | L-Dab |
| | B-LR | LS992241.1 | Polymyxin E | D | V | G | E | I | S | A | I | D | K | L-Dab |
| | PKB1 | JN660148.1 | D-Dab3-Polymyxin B | D | V | G | E | I | S | A | I | D | K | L-Dab |
| | | *Region 5* | | | | | | | | | | | | |
| **pmxE5** | Polymyxin Query | | | D | V | G | E | I | S | A | I | D | K | L-Dab |
| | E681 | EU371992.1 | Polymyxin A | D | V | G | E | I | S | A | I | D | K | L-Dab |
| | M-1 | HE577054.1 | Polymyxin P | D | V | G | E | I | S | A | I | D | K | L-Dab |
| | | KP262070.1 | Polymyxin E | D | V | G | E | I | S | A | I | D | K | L-Dab |

| | | | | | | | | | | | | | | |
|---|---|---|---|---|---|---|---|---|---|---|---|---|---|---|
| | B-LR | LS992241.1 | Polymyxin E | D | V | G | E | I | S | A | I | D | K | L-Dab |
| | PKB1 | JN660148.1 | D-Dab3-Polymyxin B | D | V | G | E | I | S | A | I | D | K | L-Dab |
| *Region 6* | | | | | | | | | | | | | | |
| **pmxA1** | Polymyxin Query | | | D | A | W | I | V | G | A | I | V | K | D-Leu |
| | E681 | EU371992.1 | Polymyxin A | D | A | W | I | V | G | A | I | V | K | D-Leu |
| | M-1 | HE577054.1 | Polymyxin P | D | A | W | T | I | A | A | I | A | K | D-Phe |
| | | KP262070.1 | Polymyxin E | D | A | W | I | V | G | A | I | V | K | D-Leu |
| | B-LR | LS992241.1 | Polymyxin E | D | A | W | I | V | G | A | I | V | K | D-Leu |
| | PKB1 | JN660148.1 | D-Dab3-Polymyxin B | D | A | W | T | I | A | A | I | A | K | D-Phe |
| *Region 7* | | | | | | | | | | | | | | |
| **pmxA2** | Polymyxin Query | | | D | F | W | N | I | G | M | V | H | K | L-Thr |
| | E681 | EU371992.1 | Polymyxin A | D | F | W | N | I | G | M | V | H | K | L-Thr |
| | M-1 | HE577054.1 | Polymyxin P | D | F | W | N | I | G | M | V | H | K | L-Thr |
| | | KP262070.1 | Polymyxin E | D | G | F | F | I | G | V | V | Y | K | L-Leu/Ile |
| | B-LR | LS992241.1 | Polymyxin E | D | G | F | F | L | G | V | V | Y | K | L-Leu/Ile |
| | PKB1 | JN660148.1 | D-Dab3-Polymyxin B | D | G | F | L | L | G | L | V | Y | K | L-Leu |
| *Region 8* | | | | | | | | | | | | | | |
| **pmxA3** | Polymyxin Query | | | D | V | G | E | I | S | A | I | D | K | L-Dab |
| | E681 | EU371992.1 | Polymyxin A | D | V | G | E | I | S | A | I | D | K | L-Dab |
| | M-1 | HE577054.1 | Polymyxin P | D | V | G | E | I | S | A | I | D | K | L-Dab |
| | | KP262070.1 | Polymyxin E | D | V | G | E | I | S | A | I | D | K | L-Dab |
| | B-LR | LS992241.1 | Polymyxin E | D | V | G | E | I | S | A | I | D | K | L-Dab |
| | PKB1 | JN660148.1 | D-Dab3-Polymyxin B | D | V | G | E | I | S | A | I | D | K | L-Dab |
| *Region 9* | | | | | | | | | | | | | | |
| **pmxA4** | Polymyxin Query | | | D | V | G | E | I | S | A | I | D | K | L-Dab |
| | E681 | EU371992.1 | Polymyxin A | D | V | G | E | I | S | A | I | D | K | L-Dab |
| | M-1 | HE577054.1 | Polymyxin P | D | V | G | E | I | S | A | I | D | K | L-Dab |
| | | KP262070.1 | Polymyxin E | D | V | G | E | I | S | A | I | D | K | L-Dab |
| | B-LR | LS992241.1 | Polymyxin E | D | V | G | E | I | S | A | I | D | K | L-Dab |
| | PKB1 | JN660148.1 | D-Dab3-Polymyxin B | D | V | G | E | I | S | A | I | D | K | L-Dab |
| *Region 10* | | | | | | | | | | | | | | |
| **pmxB1** | Polymyxin Query | | | D | F | W | N | I | G | M | V | H | K | L-Thr |
| | E681 | EU371992.1 | Polymyxin A | D | F | W | N | I | G | M | V | H | K | L-Thr |
| | M-1 | HE577054.1 | Polymyxin P | D | F | W | N | I | G | M | V | H | K | L-Thr |
| | | KP262070.1 | Polymyxin E | D | F | W | N | I | G | M | V | H | K | L-Thr |
| | B-LR | LS992241.1 | Polymyxin E | D | F | W | N | I | G | M | V | H | K | L-Thr |
| | PKB1 | JN660148.1 | D-Dab3-Polymyxin B | D | F | W | N | I | G | M | V | H | K | L-Thr |

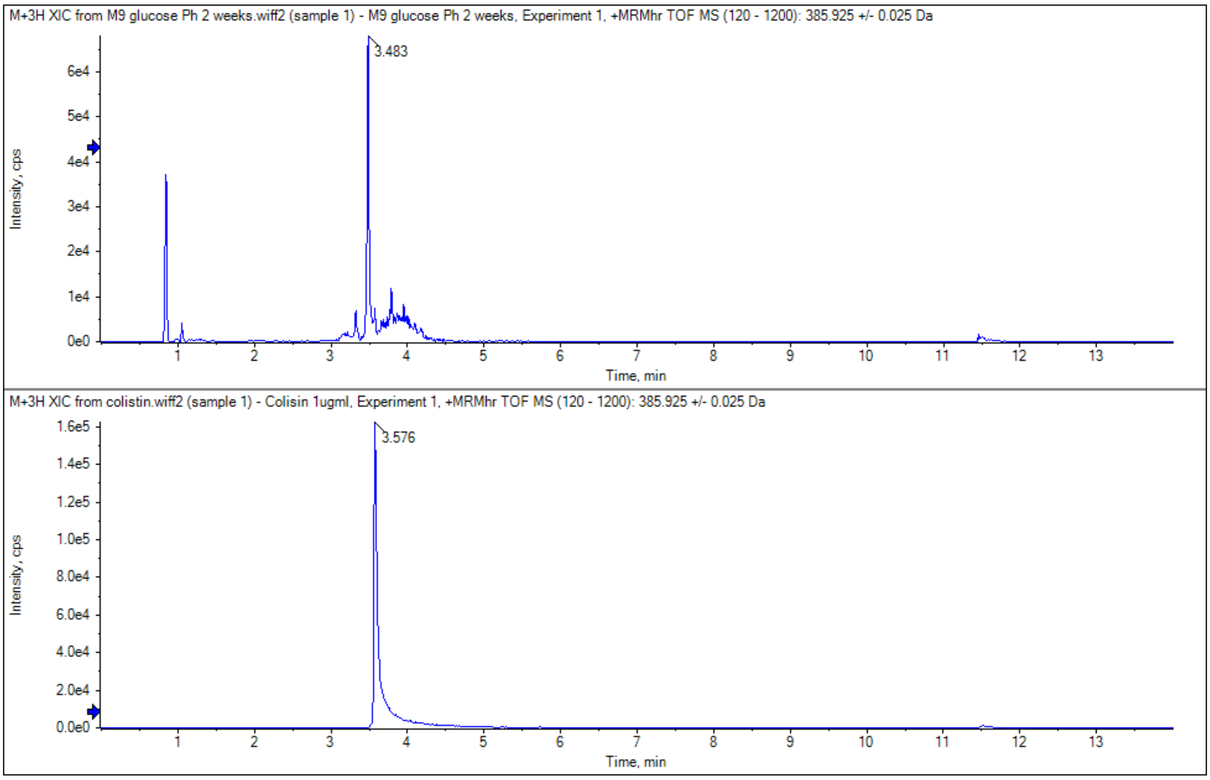


**Supplementary Figure 1** showing the chromatogram of the *P. polymyxa* 1G CFS (top) and the equivalent chromatogram of a colistin sulphate commercial sample (bottom). This highlights that they both elute under the same conditions.

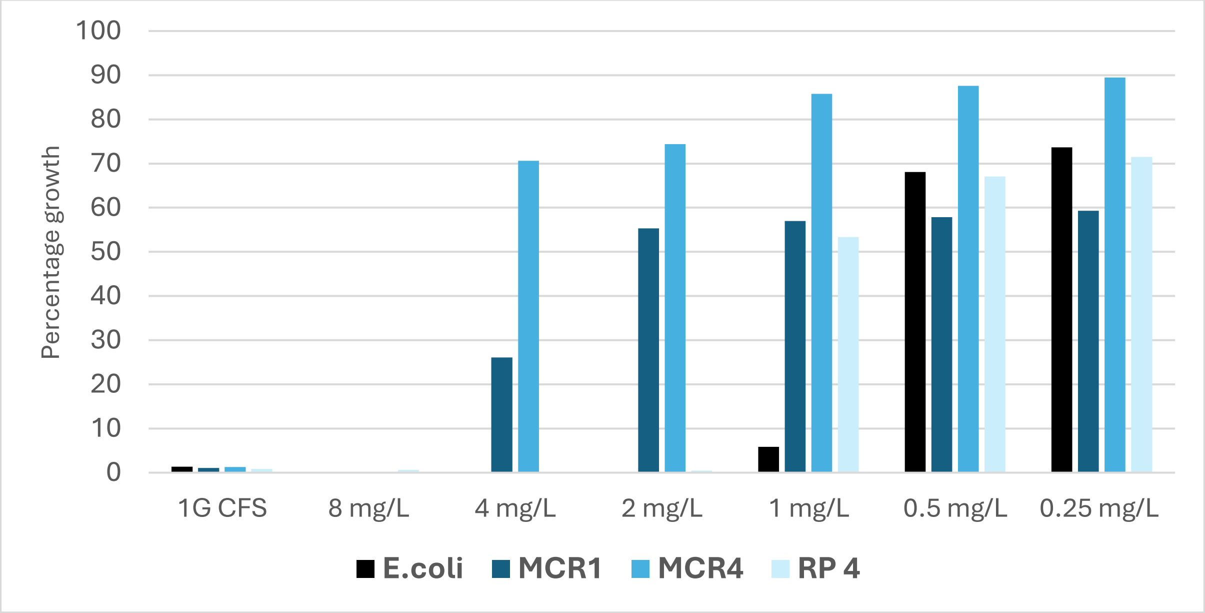


**Supplementary Figure 2:** Comparative percentage growth of resistance panel strains in the presence of colistin. This shows that the isolates containing *mcr-1* and *mcr-4* can grow in the presence of 4 mg/L colistin, above the EUCAST clinical breakpoint of 2 mg/L colistin. Isolate 24757-2026 could grow at 1 mg/L colistin, indicating a weak resistance mechanism may be present, but is not clinically resistant.
